# Earliest observation of the tetracycline destructase *tet(X3)*

**DOI:** 10.1101/2023.08.29.555419

**Authors:** Frédéric Grenier, Simon Lévesque, Sébastien Rodrigue, Louis-Patrick Haraoui

## Abstract

Tigecycline is an antibiotic of last resort for infections with carbapenem-resistant *Acinetobacter baumannii*. Plasmids harboring variants of the tetracycline destructase *tet(X)* promote rising tigecycline resistance rates. We report the earliest observation of *tet(X3)* in a clinical strain predating tigecycline’s commercialization, suggesting selective pressures other than tigecycline contributed to its emergence.

## Introduction

Antimicrobial resistance (AMR) is a major global public health issue, with AMR deaths surpassing HIV and malaria in terms of infectious disease-related mortality (1). In 2018, the World Health Organization (WHO) issued its priority list for discovery, research, and development of new treatments for antibiotic-resistant bacteria. All priority 1 critical pathogens – *Acinetobacter baumannii, Pseudomonas aeruginosa* and *Enterobacteriaceae* – share a common trait: resistance to carbapenems, a class within the broad beta-lactam family of antibiotics (2).

Therapeutic options for carbapenem-resistant *Acinetobacter baumannii* are often limited to tigecycline and colistin. Resistance to these antibiotics is increasing, primarily as a result of variants of *tet(X)* and *mcr* genes respectively. *tet(X3)*, the predominant *tet(X)* variant among *Acinetobacter* spp., was initially reported in 2019 in an *A. baumannii* isolated in China in 2017 (3). Retrospective analyses have since highlighted the role of non-*baumannii Acinetobacter* in the global distribution of *tet(X3)*, with the earliest clinical isolates dating back to 2010 based on a search in GenBank. In this study, we report an *A. junii* strain (Ajun-H1-2) harboring *tet(X3)*, isolated from a blood culture in 2004 in Israel.

## Materials and Methods

### Strains

We obtained 198 clinical *Acinetobacter* spp. isolated in Israel between 2001 and 2006 from three archives: 1) directly from Chaim Sheba Medical Center, Tel Hashomer, Israel (n= 140); 2) from JMI Laboratories who, as part of their SENTRY Antimicrobial Surveillance Program, still held 37 isolates also from Chaim Sheba Medical Center, distinct from those we obtained directly from the medical center; and 3) 21 isolates from International Health Management Associates inc. (IHMA), collected from two different anonymized hospitals.

### Whole Genome Sequencing

All isolates were grown in lysogenic broth at 37°C overnight. DNA libraries were prepared from extracted gDNA using the NEBNext Ultra II FS DNA Library Prep Kit for Illumina (NEB). DNA was purified and size selected using Ampure XP beads (Beckman Coulter) and quantified using Quant-it PicoGreen dsDNA assay (Thermo Fisher). The quality and size distribution of the libraries were assessed on a Fragment Analyzer using the HS NGS Fragment Kit (Agilent). The pooled samples were then sequenced on a NovaSeq6000 (Illumina) with 250 bp paired-end sequencing.

A subset of strains, including Ajun-H1-2, was also sequenced using an Oxford Nanopore Technologies (ONT) R10.4 Flow Cell on a MinION Mk1B. Extracted gDNA was treated with the NEBNext Ultra II End Repair/dA-Tailing Module (NEB). Barcodes from the Native Barcoding Expansion 1-12 & 13-24 from ONT were ligated using the NEBNext Ultra II Ligation Module (NEB). DNA was purified using Ampure XP beads (Beckman Coulter). The DNA from different barcoded samples was pooled and the adapter AMII (ONT) was ligated using the NEBNext Ultra II Ligation Module (NEB).

### Data Analysis and Bioinformatics

For Illumina reads, quality assessment and trimming were done using fastp 0.21.0 with --cut_right --cut_window_size 4 --cut_mean_quality 20 --length_required 30 -- detect_adapter_for_pe (4). Assemblies were made using Unicycler 0.4.9 (5) with the trimmed Illumina short reads and ONT long reads when available. Contigs were filtered to retain only those above 500 bp.

Taxonomic identification was made on the assemblies using Kraken 2 (2.0.9-beta) (6). Antibiotic resistance genes (ARGs) were searched using ResFinder 4.1 (7). Detected *bla*_OXA_ variants were curated using BLDB (8). Assemblies were annotated with Prokka 1.14.5 (9) using the additional databases Pfam, TIGRFAM, and the *bla*_OXA_ variants present in BLDB (8). The plasmid figure was generated using AliTV (10).

### Antibiotic susceptibility testing

Antibiotic susceptibility testing of Ajun-H1-2 was carried out using the AST-N801 card on a Vitek2 (bioMérieux) and the Sensititre GNX3F plate (ThermoFisher).

### Data Availability

Sequencing reads are deposited in GenBank under BioProject number PRJNA845765.

## Results

Strain Ajun-H1-2, identified as *A. junii* based on molecular analyses, was isolated from a blood culture in 2004 in Israel. No further information about the patient from which it was isolated is available. The strain carried numerous ARGs, including both the *tet(X3)* tetracycline destructase and the *bla*_OXA-58_ carbapenemase (see Table 1). All ARGs were located on a 367 kilobase (kb) plasmid named pTet(X3)-Ajun-H1-2. No origin of replication was identified. Strain Ajun-H1-2 also contained 4 other small plasmids ranging in size between 2 and 8 kb.

**Table 1.** **a)** Antibiotic-resistance genes found in Ajun-H1-2 were all present on pTet(X3)-Ajun-H1-2. **b)** Antibiotic susceptibility results obtained using the AST-N801 card on a Vitek2 and a Senstitre GNX3F plate. MIC: minimal inhibitory concentration.

| Antibiotic Family | Resistance gene | Identity | Query / Template length | Position in contig | Predicted phenotype | Accession number |
| --- | --- | --- | --- | --- | --- | --- |
| aminoglycoside | aadA1 | 100 | 789 / 789 | 284767..285555 | Aminoglycoside resistance | JQ480156 |
| aminoglycoside | ant(2'')-Ia | 100 | 534 / 534 | 231965..232498 | Aminoglycoside resistance<br>Alternate name; ant(2'')-Ia | X04555 |
| aminoglycoside | aph(3')-Ia | 100 | 816 / 816 | 4518..5333 | Aminoglycoside resistance | X62115 |
| beta-lactam | blaOXA-58 | 100 | 843 / 843 | 536..1378 | Beta-lactam resistance | AY665723 |
| macrolide | mph(E) | 100 | 885 / 885 | 355941..356825 | Macrolide resistance | DQ839391 |
| macrolide | msr(E) | 100 | 1476 / 1476 | 356881..358356 | Macrolide, Lincosamide and Streptogramin B resistance | FR751518 |
| phenicol | floR | 98.35 | 1214 / 1215 | 9936..11149 | Phenicol resistance | AF118107 |
| sulphonamide | sul2 | 100 | 816 / 816 | 27860..28675 | Sulphonamide resistance | AY034138 |
| tetracycline | tet(X3) | 100 | 1167 / 1167 | 32051..33217 | Tetracycline Resistance | MK134375 |
| trimethoprim | dfrA1 | 100 | 474 / 474 | 283617..284090 | Trimethoprim resistance | X00926 |

**Table 1 a)** Antibiotic-resistance genes found in Ajun-H1-2 were all present on pTet(X3)-Ajun-H1-2. **b)** Antibiotic susceptibility results obtained using the AST-N801 card on a Vitek2 and a Sensititre GNX3F plate. MIC: minimal inhibitory concentration.
| Antibiotic | Vitek MIC | Vitek Interpretation | Sensititre MIC | Sensititre Interpretation |
| --- | --- | --- | --- | --- |
| Tetracycline | ≥ 16 | R |  |  |
| Doxycycline |  |  | 4 | S |
| Minocycline |  |  | 2 | S |
| Tigecycline | 1 | S | 2 | S |
| Ampicillin/sulbactam | ≤ 2 | S | 4/2 | S |
| Ticarcillin/calvulanic acid |  |  | 16/2 | S |
| Piperacillin/tazobactam | ≤ 4 | S | 8/4 | S |
| Ceftazidime | ≤ 1 | S | 2 | S |
| Ceftolazone/tazobactam | ≤ 0.25 | S |  |  |
| Cefepime | 0.5 | S | 2 | S |
| Imipenem | ≤ 0.25 | S | 1 | S |
| Meropenem | ≤ 0.25 | S | 1 | S |
| Doripenem |  |  | 0.5 | S |
| Ciprofloxacin | 2 | I | 2 | I |
| Levofloxacin | 2 | S | 1 | S |
| Amikacin | ≤ 2 | S | 4 | S |
| Gentamicin | ≥ 16 | R | 8 | I |
| Tobramycin | ≥ 16 | R | 8 | I |
| Trimethoprim/sulfamethoxazole | ≥ 320 | R | 4/76 | R |

Based on literature and GenBank searches, 12 other cases of *tet(X3)* and *bla*_OXA-58_ co-existence on the same plasmid have been reported among more recently isolated *Acinetobacter* spp. from healthcare settings in the United States (7 *A. baumannii*) and Pakistan (1 *A. junii*), as well as from farm animals in China (3 *A. towneri* and 1 *Acinetobacter* spp.) (11–13). None of these 12 plasmids resembled pTet(X3)-Ajun-H1-2 (Figure 1).

**Figure 1:**
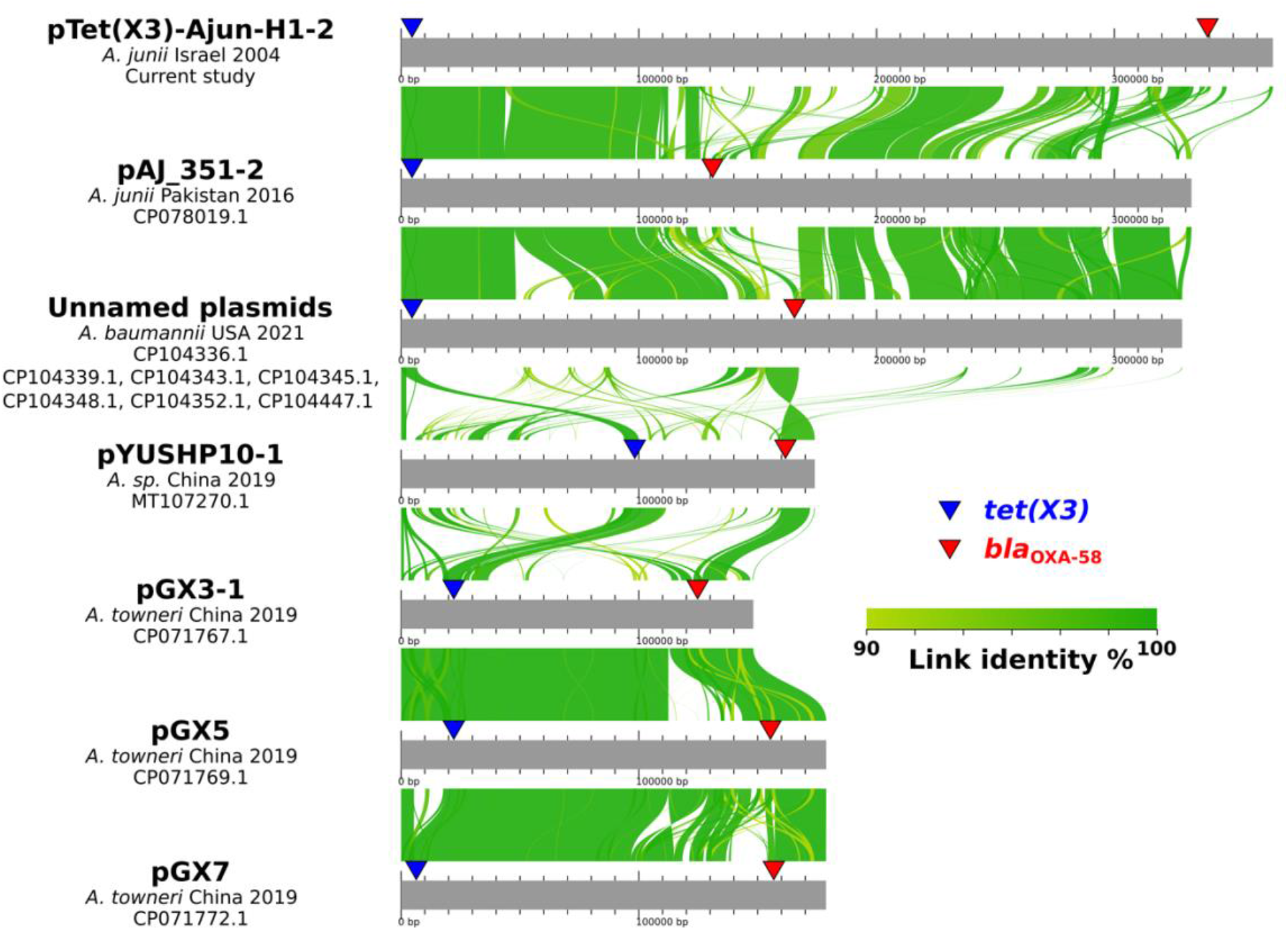
Alignments of plasmids harboring both *tet(X3)* and *bla*_OXA-58_ including pTet(X3)-Ajun-H1-2 reported in this study. Alignments done using AliTV (10).

Susceptibility testing with 19 different antibiotics was performed on Ajun-H1-2, demonstrating decreased susceptibility (intermediate or resistant) to ciprofloxacin, colistin, gentamicin, tetracycline, tobramycin, and trimethoprim/sulfamethoxazole with varying minimal inhibitory concentrations (MICs) depending on the method used. The strain was sensitive to other antibiotics tested, including all beta-lactams, doxycycline, minocycline and tigecycline (see Table 1).

## Discussion

We report the earliest evidence of *tet(X3)* in an *A. junii* strain from 2004. This gene is capable of conferring resistance to most members of the tetracycline family of antibiotics, including tigecycline, initially commercialized in 2005. Despite this, Ajun-H1-2 was sensitive to minocycline, doxycycline and tigecycline, though resistant to tetracycline. Such discrepancies between *tet(X3)* presence and resistance profile have been primarily noted in strains isolated in farm animals. In one report, 46/47 *tet(X3)*-positive *Acinetobacter* strains were sensitive to tigecycline, with the remaining strain, an *A. indicus*, demonstrating an intermediate susceptibility. Among these same 47 strains, only 4 were found to be resistant to doxycycline and to minocycline (tetracycline was not assessed). Such discrepancies could be related to host factors modulating susceptibility to the tetracycline family of antibiotics, to gene promoters upstream of *tet(X3)* regulating expression levels, or to *tet(X3)* variants present and their copy numbers.

Although cases of *tet(X3)*-positive *Acinetobacter* spp. have been isolated on all continents, most reports come from China, which may reflect sampling and reporting biases. Tetracycline antibiotics are widely used in various settings in China, including in animal husbandry and other agricultural practices (14), which could also explain a greater number of reported cases.

In a recent large-scale survey of *tet(X)*-positive bacteria in China among humans, animals and environmental niches, about one third of *tet(X)*-positive bacteria carried the *tet(X3)* variant, nearly equal to the most common variant, *tet(X2)* (15). Whereas all *tet(X3)*-positive bacteria belonged to the *Acinetobacter* genus, with a predominance of *A. indicus*, the *tet(X2)*-positive bacteria were found in numerous genera. A few *Acinetobacter* isolates carried *tet(X5)* and *tet(X6)* variants, the latter not limited to *Acinetobacter* spp., and some *Acinetobacter* isolates carried both *tet(X3)* and *tet(X6)*.

In conclusion, our report further supports the role of non-*baumannii Acinetobacter* in the initial dissemination of *tet(X3)*. Whereas the use of tigecycline has been linked to the rise of *tet(X)* variants, this study demonstrates that *tet(X3)* predated the commercialization of this antibiotic. Finally, our study’s findings highlight the limitations of relying on antibiotic susceptibility testing as a means of retrospectively tracking the emergence and spread of ARGs. Further research is needed to more fully understand the origins of *tet(x3)* as well as minimal inhibitory concentration variations among *tet(x3)*-positive strains.

## Acknowledgments

The authors would like to thank Gill Smollan and Sharon Amit at Chaim Sheba Medical Center, Tel HaShomer, Ramat Gan, Israel.

## Disclaimers

The authors have no conflicts of interest to declare.

